# LoFTK: a framework for fully automated calculation of predicted Loss-of-Function variants

**DOI:** 10.1101/2021.08.09.455694

**Authors:** A. Alasiri, K. J. Karczewski, B. Cole, B. Loza, J. H. Moore, S. W. van der Laan, F. W. Asselbergs, B. J. Keating, J. van Setten

## Abstract

**Motivation:** Loss-of-Function (LoF) variants in human genes are important due to their impact on clinical phenotypes and frequent occurrence in the genomes of healthy individuals. Current approaches predict high-confidence LoF variants without identifying the specific genes or the number of copies they affect. Moreover, there is a lack of methods for detecting knockout genes caused by compound heterozygous (CH) LoF variants.

**Results:** We have developed the Loss-of-Function ToolKit (LoFTK), which allows efficient and automated prediction of LoF variants from both genotyped and sequenced genomes. LoFTK enables the identification of genes that are inactive in one or two copies and provides summary statistics for downstream analyses. LoFTK can identify CH LoF variants, which result in LoF genes with two copies lost. Using data from parents and offspring we show that 96% of CH LoF genes predicted by LoFTK in the offspring have the respective alleles donated by each parent.

**Availability and implementation:** LoFTK is an open source software and is freely available to non-commercial users at https://github.com/CirculatoryHealth/LoFTK

**Supplementary information:** Supplementary data are available at *Bioinformatics* online.

## 1. Introduction

Loss-of-function (LoF) variants are determined to have a critical effect on gene function by inactivating protein-coding genes (MacArthur and Tyler-Smith 2010). Remarkably, recent analyses of the human genome have uncovered that individuals harbor many dozens of LoF variants, including stop-gained, frameshift variants and splice site disruptions (Balasubramanian et al. 2011; MacArthur et al. 2012). On average, LoF variants are deleterious, and thus usually tend to be found at very low frequencies in the human population. These variants can have a profound impact on the gene transcripts and translated proteins. The association of LoF variants with complex diseases and phenotypic traits may lead to the discovery and validation of novel therapeutic targets (MacArthur et al. 2012). However, hundreds of LoF variants were found to be functionally neutral with no detectable influence on phenotypes (1000 Genomes Project Consortium et al. 2012; Lek et al. 2016).

Several difficulties emerge when evaluating LoFs on a broad scale. False positives in the prediction of LoF variants can arise due to artifacts that may occur during genotype calling, mapping, and annotation (MacArthur et al. 2012). To annotate high-confidence (HC) LoF variants only, Loss-Of-Function Transcript Effect Estimator (LOFTEE) (Karczewski et al. 2020) can be used. LOFTEE is a plugin implemented in the Ensembl Variant Effect Predictor (VEP) (McLaren et al. 2016) that imposes stringent filtering criteria to annotate HC LoF variants, eliminate nonsense mutations that are unlikely to impact protein function, and exclude LoF variants that are enriched with annotation artifacts. However, LoF variants discovery can also be used to predict single-copy losses (heterozygous LoF variants) that inactivate a single copy of a gene, or two-copy losses that completely knockout a gene. Two-copy losses can be caused by homozygous and compound heterozygous (CH) LoF variants. CH variants appear when parents both donate a LoF-causing allele that locates at different loci in the same gene (Kamphans et al. 2013). There is mounting evidence that CH LoF variants have a role in complex diseases. For example, both homozygous and CH LoF variants have been found to increase the risk of autism spectrum disorder (Yu et al. 2013; Lim et al. 2013). Current tools, such as LOFTEE, merely annotate HC LoF variants and generate a standard VEP output, but do not distinguish between single-copy LoF genes and two-copy LoF genes.

Here we present an open source tool, the Loss-of-Function ToolKit (LoFTK), which allows efficient and automated prediction of LoF variants from genotyped and sequenced genomes. LoFTK analyzes genetic data in three steps; 1) LoF annotation, 2) prediction of one-copy loss and two-copy loss of genes, and 3) generation of a summary statistics report describing the total number of LoF variants (homozygous, heterozygous and CH), LoF genes (single-copy and two-copy loss), and the average, minimum and maximum numbers of LoF variants and genes per sample.

## 2. Materials and methods

### 2.1. Main workflow

The LoFTK workflow consists of 4 analytical steps visualized in Figure 1 and described below.

**Figure 1:**
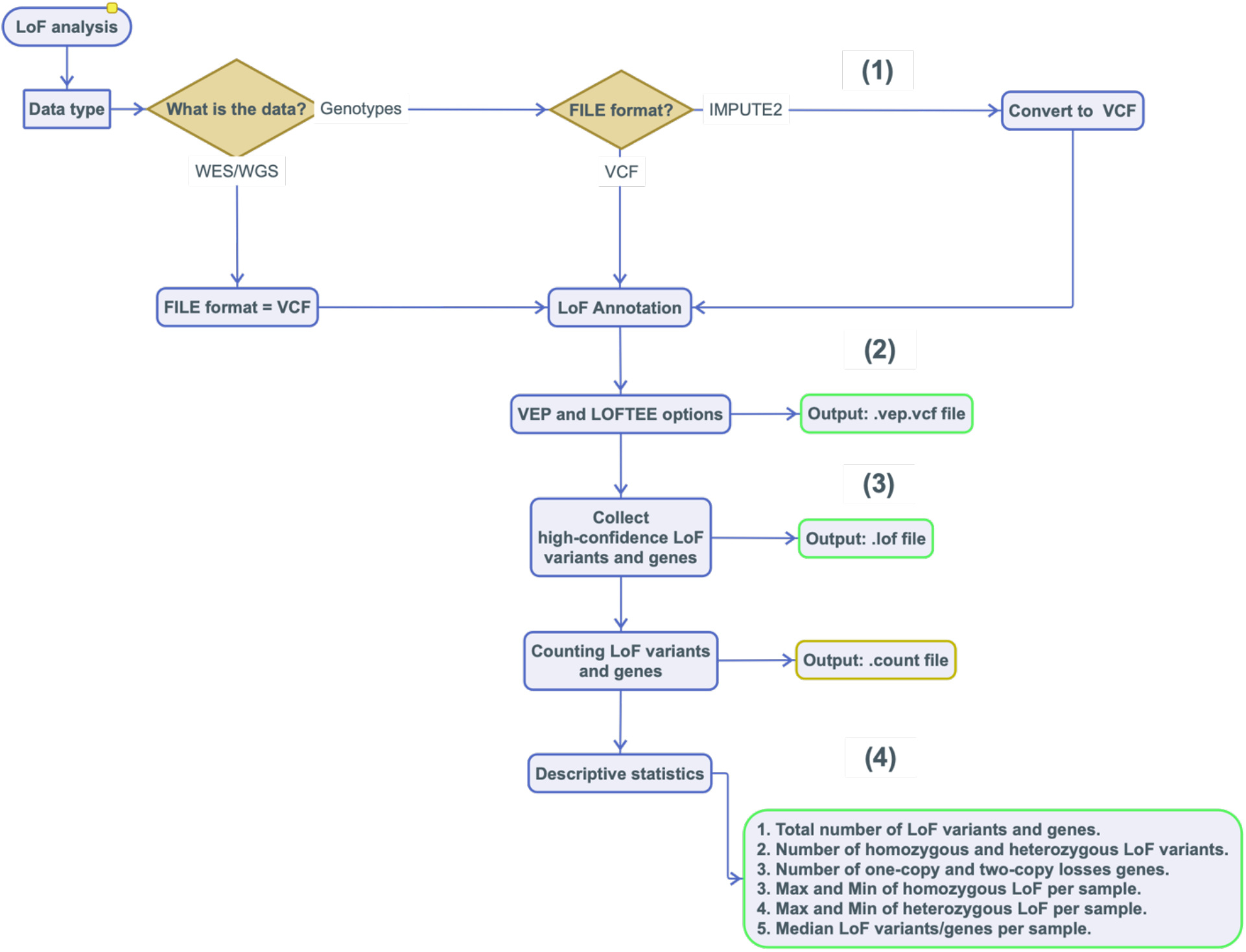
The workflow of LoFTK pipeline. Four steps involved in LoFTK; (1) preprocessing from IMPUTE2 to VCF, (2) LoF annotation and vep.vcf.file creation, (3) filtering HC LoF variants and counting LoF variants and genes, and (4) descriptive analysis.

#### 2.1.1. Preprocessing: conversion of IMPUTE2 to VCF

The first step depends on the input data formats. Two common file formats are permitted as inputs; IMPUTE2 (Howie et al. 2012; Howie et al. 2009) output format and the Variant Call Format (VCF). The input data has to contain phased genotypes for distinguishing compound heterozygotes from two variants on the same allele. LoFTK uses two quality metrics for imputed genotypes: the imputation quality (info score) and imputed allele probability. The imputation quality contains values between 0 and 1, where higher values mean that a variant has been imputed with more certainty. Besides, imputation methods generate a probabilistic prediction of the missing genotypes, which stands for the likelihood of carrying genotypes combinations of A/A, A/B, and B/B in a particular individual. The supreme estimated genotype is the genotype that has the highest likelihood of being correct (Marchini and Howie 2010). LoFTK has cut-off options to filter based on the optimal imputation quality metrics (Supplementary Material). After filtering, IMPUTE2 files are converted to VCF files. The VCF files that are generated from IMPUTE2 files or introduced directly by the user are applied as an input to the next step.

#### 2.1.3. LoF annotation

The second step consists of annotation of LoF variants using VEP and LOFTEE. LOFTEE utilizes the Ensembl API framework to annotate HC LoF variants. We designed LoFTK to be capable of processing data with *Homo sapiens* (human) genome assemblies GRCh37 and GRCh38, and it can easily be upgraded to future genome builds. The VEP will return results as VEP VCF format, which is similar to the input VCF, but in addition shows LoF information in the INFO field, such as LoF flags and LoF filtering outcome (high-confidence or low-confidence) and LoF flags.

#### 2.1.4. Calculation of LoF variants and genes

From the VEP output, the HC LoF variants are filtered, followed by parallel determination of homozygous and heterozygous LoF variants (Table 2) and allele frequencies, as well as the copy number loss (single-copy or two-copy) of LoF genes (Table 1). LoFTK recognizes CH LoF variants, which result in LoF genes with two copies losses. We used imputed genotypes from the Genome of the Netherlands (GoNL) project as family-based trio data to confirm both parents donating a single LoF allele to proband at distinct loci in the same gene (Supplementary Material). Optionally, LoFTK can be used to determine ‘mismatched genes’ between samples; these are genes that are active in one or two copies in one sample and completely inactive in the other sample. This feature helps study interactions between human genomes, for instance during pregnancy (maternal vs fetal genome) and after stem cell or solid organ transplantation (donor vs recipient genome).

**Table 1:** The output of predicted LoF genes from WES in UKBB. This table shows the predicted LoF gene ID and symbol in column 1 and 2, respectively. The third column represents the frequency of single-copy loss gene, while the fourth represents the frequency of two-copy losses gene. The rest of columns indicate the number of copy losses for each individual; 0 for not carrying LoF gene, 1 for sigle-copy LoF gene and 2 for two-copy LoF genes.

| gene_ID | gene_symbol | single_copy_LoF_frequency | two_copy_LoF_frequency | Sample 1 | Sample 2 | Sample 3 |
| --- | --- | --- | --- | --- | --- | --- |
| ENSG00000198464 | ZNF480 | 0.387846291 | 0.535299374 | 2 | 1 | 1 |
| ENSG00000105877 | DNAH11 | 0.502234138 | 0.250893655 | 1 | 0 | 1 |
| ENSG00000152936 | IFLTD1 | 0.013181412 | 0 | 0 | 1 | 0 |
| ENSG00000039537 | C6 | 0.000446828 | 0.000223414 | 0 | 0 | 0 |
| ENSG00000221938 | OR2A14 | 0 | 0.000223414 | 0 | 0 | 0 |

**Table 2:** The output of predicted LoF variants from WES in UKBB. high-confidence LoF variants are listed in the first column, followed by their consequences in the second column. The third and fourth columns show frequencies of heterozygous and homozygous LoF variants, respectively. The rest of columns indicate the zygosity of LoF variants for each individual; 0 for not carrying LoF variant, 1 for hetrozygous LoF variant and 2 for homozygous LoF variant.

| SNP_ID | Consequence | single_copy_LoF_frequency | two_copy_LoF_frequency | Sample 1 | Sample 2 | Sample 3 |
| --- | --- | --- | --- | --- | --- | --- |
| chr19_52300416_CTG_C | frameshift_variant | 0.385570391 | 0.538972204 | 1 | 1 | 2 |
| chr7_21543345_G_T | stop_gained | 0.498856177 | 0.250090958 | 0 | 1 | 1 |
| chr10_72508273_T_C | splice_acceptor_variant | 0.425123229 | 0.449918512 | 2 | 1 | 0 |
| chr1_3602477_AC_A | frameshift_variant | 4.98E-06 | 0 | 1 | 0 | 0 |
| chr7_21543345_G_T | stop_gained | 0.498856177 | 0.250090958 | 0 | 2 | 1 |

#### 2.1.5. Descriptive analysis

Finally, descriptive statistics of LoF variants are calculated, such as the total number of LoF variants, number of single-copy and two-copy LoF genes, and median of LoF variants per participant.

### 2.2. Imputation quality threshold

The imputed genotype data provides two quality metrics: the INFO score and the imputed alleles probability. UK Biobank (UKBB) was used as the gold standard for determining the optimal quality metrics for obtaining the most genuine LoF variants from imputed genotypes data. We retrieved whole exome sequencing (WES) and array genotypes data from 4,476 randomly selected UKBB participants. Both data were phased using SHAPEIT2 (Howie et al. 2009; Howie et al. 2012) and array genotypes were imputed by IMPUTE2 (Delaneau et al. 2011). A combined reference panel from the 1000 Genome project phase 3 (1000 Genomes Project Consortium et al. 2015) and Genome of the Netherlands (GoNL) study (Boomsma et al. 2014) was used for phasing and imputation. We used LoFTK for LoF analysis in phased WES and three datasets of variants in imputed genotypes data. These subsets were divided based on variants with INFO scores above: 0.3, 0.6, 0.9. for each individual, predicted LoF variants in the WES were compared to LoF variants in each subset with considering the imputed allele probabilities ranging from 0.01 to 0.1 for that variant (Supplementary Figure 1), in order to count the number of false negatives (average of LoF variants predicted in WES data but not in imputed data) and false positives (average of LoF variants predicted in imputed data but not in WES data).

### 2.3. Validity of predicted CH LoF in trios

LoFTK is capable of annotating CH LoF variants, which introduce two inactive copies of a gene. To confirm the transmission of genuine CH LoF variants from parents to probands, we used trio-family genotype data from the Genome of the Netherlands (GoNL) cohort (Illumina Immunochip microarray SNP data) (Boomsma et al. 2014). We performed a quality control step as preprocessing filtrations to impute genotypes data (Supplementary Material). We used the TOPMed imputation server (Taliun et al. 2021) to impute untyped variants in 760 individuals from 250 families. LoFTK predicted LoF variants from imputed genotypes and we investigated transmission of CH LoF variants from parents to offspring.

## 3. Results

### 3.1. LoFTK software

Perl and bash were used to develop LoFTK for identifying LoF variants. Instructions on how to install and run LoFTK as well as example datasets are publicly available at https://github.com/CirculatoryHealth/LoFTK. LoFTK is highly customizable with options and directories settings, and all the options are explained in the LoF.config file and GitHub README. It is designed to run as a command line program with user-friendly flags, which helps non-experts users to get quickly familiarized. LoFTK requires pre-installed tools, such as BASH, Perl (>= version 5.10.1), Ensembl VEP (https://github.com/Ensembl/ensembl-vep) and LOFTEE (https://github.com/konradjk/loftee). We tested LoFTK on CentOS 7 managed by SLURM or SGE.

### 3.2. Generation of LoF variants and genes

LoFTK uses the information present in large-scale sequencing and genotyping data to generate two matrices of LoF variants and their respective genes, a list of LoF variants allele frequencies, and a report with descriptive statistics on the variants and genes. In the LoF variants matrix, the variants are represented as rows, and individuals are represented as columns. Each matrix’s cell contains a number that represents the homozygous or heterozygous status of a given LoF variant for a given individual as shown in Table 1. Similarly, the columns in the LoF genes matrix define individuals except the rows represent the LoF genes, and each number in the matrix cell indicates that either the gene has no copy loss (0), single-copy loss (1) or two-copy loss (2) (Table 2). Finally, LoFTK generates information file with “.info” extension to show descriptive statistical report for predicted LoF variants and genes, such as the total LoF variants and genes, total heterozygous and homozygous LoF variants, total single-copy and two-copy LoF genes, and median of LoF variants and genes per participant.

### 3.3. Imputation quality cut-off points

We assessed imputation quality metrics for obtaining the most genuine LoF variants in imputed genotype data by comparing existence of each predicted LoF variant between WES and three imputed dataset (INFO > 0.3, 0.6, 0.9) with considering the imputed allele probability cutoffs between 0.01 to 0.1 (see Section 2.2.).

LoFTK analysis for imputed dataset with INFO > 0.9 shows an optimal prediction of true LoF variants, because it has less false positive 2-copy LoF variants compared to the others (0.3 and 0.6) (Supplementary Figure 1). However, choosing an optimal imputed allele probability was difficult due to the lack of apparent variations.

### 3.4. CH LOF variants in trios

CH LoF variants occur when both parents donate a single LoF allele to proband at distinct loci within the same gene. We used trio-families from the GoNL to evaluate the accuracy of obtaining two inactive copies in genes caused by CH LoF variants (see Section 2.3.).

We predicted LoF variants and genes in 250 families’ imputed genotypes (760 individuals). We found 642 LoF variants affecting 571 genes (Table 3). In 164 probands, we identified 250 events of CH LoF variants producing 2-copy LoF genes. There were 240 (96%) true transmissions of CH LoF in parent-offspring, whereas there were 10 false transmissions.

**Table 3:** Predicted LoF variants and genes in the GoNL. LoF genes column shows numbers of total LoF genes, one copy inactive genes (1-copy) and two copies inactive genes (2-copy). LoF variants shows total number of predicted LoF variants, heterozygous and homozygous variants.

|  | *LoF genes* | *LoF variants* |
| --- | --- | --- |
| *Total* | 571 | 642 |
| *1-copy* | 571 | - |
| *2-copy* | 196 | - |
| *Median 1-copy per individual* | 49 | - |
| *Median 2-copy per individual* | 21 | - |
| *Heterozygous* | - | 641 |
| *Homozygous* | - | 213 |
| *Median heterozygous per individual* | - | 54 |
| *Median homozygous per individual* | - | 21 |

## 4. Conclusions

Prediction of LoF variants and genes provide important insight into the discovery of possible disease-causing mutations and potential therapeutic targets. LoFTK is easy to use and helps users to predict LoF variants from genotyped and sequenced genomes, identifying genes that are inactive in 1 or 2 copies, and providing summary statistics report describing the total number of LoF variants, LoF genes, and their average, minimum and maximum per sample. LoFTK is highly customizable and extra features for the identification of knockout genes in copy number variation (CNV) and predicting the pathogenicity of predicted LoF variants can be easy added.

## Acknowledgements

This study was carried out utilizing the UK Biobank Resource under application number [24711]. This study makes use of data generated by the Genome of the Netherlands Project. Funding for the project was provided by the Netherlands Organization for Scientific Research under award number [184021007], dated (July 9, 2009) and made available as a Rainbow Project of the Biobanking and Biomolecular Research Infrastructure Netherlands (BBMRI-NL). Samples where contributed by LifeLines (http://lifelines.nl/lifelines-research/general), The Leiden Longevity Study (http://www.healthy-ageing.nl; http://www.langleven.net), The Netherlands Twin Registry (NTR: http://www.tweelingenregister. org), The Rotterdam studies, (http://www.erasmus-epidemiology.nl/rotterdamstudy) and the Genetic Research in Isolated Populations program (http://www.epib.nl/research/geneticepi/research.html#gip). The sequencing was carried out in collaboration with the Beijing Institute for Genomics (BGI). We are thankful for the support of the ERA-CVD program ‘druggable-MI-targets’ (grant number: 01KL1802), the EU H2020 TO_AITION (grant number: 848146), and the Leducq Fondation ‘PlaqOmics’.

## Data availability statement

The data used in this article were provided by the UK Biobank under application (#24711) and the Genome of the Netherlands under application number (#2021217). Access to these data can be achieved by request from UK Biobank (https://www.ukbiobank.ac.uk/enable-your-research/apply-for-access) and the Genome of the Netherlands (https://www.nlgenome.nl/menu/main/app-request)

## References

1000 Genomes Project Consortium, Goncalo R. Abecasis, Adam Auton, Lisa D. Brooks, Mark A. DePristo, Richard M. Durbin, Robert E. Handsaker, Hyun Min Kang, Gabor T. Marth, and Gil A. McVean. 2012. “An Integrated Map of Genetic Variation from 1,092 Human Genomes.” Nature 491 (7422): 56–65.

Balasubramanian, Suganthi, Lukas Habegger, Adam Frankish, Daniel G. MacArthur, Rachel Harte, Chris Tyler-Smith, Jennifer Harrow, and Mark Gerstein. 2011. “Gene Inactivation and Its Implications for Annotation in the Era of Personal Genomics.” Genes & Development 25 (1): 1–10.

Boomsma, Dorret I., Cisca Wijmenga, Eline P. Slagboom, Morris A. Swertz, Lennart C. Karssen, Abdel Abdellaoui, Kai Ye, et al. 2014. “The Genome of the Netherlands: Design, and Project Goals.” European Journal of Human Genetics: EJHG 22 (2): 221–27.

Kamphans, Tom, Peggy Sabri, Na Zhu, Verena Heinrich, Stefan Mundlos, Peter N. Robinson, Dmitri Parkhomchuk, and Peter M. Krawitz. 2013. “Filtering for Compound Heterozygous Sequence Variants in Non-Consanguineous Pedigrees.” PloS One 8 (8): e70151.

Karczewski, Konrad J., Laurent C. Francioli, Grace Tiao, Beryl B. Cummings, Jessica Alföldi, Qingbo Wang, Ryan L. Collins, et al. 2020. “The Mutational Constraint Spectrum Quantified from Variation in 141,456 Humans.” Nature 581 (7809): 434–43.

Lek, Monkol, Konrad J. Karczewski, Eric V. Minikel, Kaitlin E. Samocha, Eric Banks, Timothy Fennell, Anne H. O’Donnell-Luria, et al. 2016. “Analysis of Protein-Coding Genetic Variation in 60,706 Humans.” Nature 536 (7616): 285–91.

Lim, Elaine T., Soumya Raychaudhuri, Stephan J. Sanders, Christine Stevens, Aniko Sabo, Daniel G. MacArthur, Benjamin M. Neale, et al. 2013. “Rare Complete Knockouts in Humans: Population Distribution and Significant Role in Autism Spectrum Disorders.” Neuron 77 (2): 235–42.

MacArthur, Daniel G., Suganthi Balasubramanian, Adam Frankish, Ni Huang, James Morris, Klaudia Walter, Luke Jostins, et al. 2012. “A Systematic Survey of Loss-of-Function Variants in Human Protein-Coding Genes.” Science 335 (6070): 823–28.

MacArthur, Daniel G., and Chris Tyler-Smith. 2010. “Loss-of-Function Variants in the Genomes of Healthy Humans.” Human Molecular Genetics 19 (R2): R125–30.

Marchini, Jonathan, and Bryan Howie. 2010. “Genotype Imputation for Genome-Wide Association Studies.” Nature Reviews. Genetics 11 (7): 499–511.

McLaren, William, Laurent Gil, Sarah E. Hunt, Harpreet Singh Riat, Graham R. S. Ritchie, Anja Thormann, Paul Flicek, and Fiona Cunningham. 2016. “The Ensembl Variant Effect Predictor.” Genome Biology 17 (1): 122.

Taliun, Daniel, Daniel N. Harris, Michael D. Kessler, Jedidiah Carlson, Zachary A. Szpiech, Raul Torres, Sarah A. Gagliano Taliun, et al. 2021. “Sequencing of 53,831 Diverse Genomes from the NHLBI TOPMed Program.” Nature 590 (7845): 290–99.

Yu, Timothy W., Maria H. Chahrour, Michael E. Coulter, Sarn Jiralerspong, Kazuko Okamura-Ikeda, Bulent Ataman, Klaus Schmitz-Abe, et al. 2013. “Using Whole-Exome Sequencing to Identify Inherited Causes of Autism.” Neuron 77 (2): 259–73.

