## Supplementary material for "LoFTK: a framework for fully automated calculation of predicted Loss-of-Function variants"

### Sections

- Determining the optimal imputation quality threshold
- Compound heterozygous LoF variants
- Supplementary Table
- Supplementary Figure

### Determining the optimal imputation quality threshold

LoFTK was developed to analyze any genetic data, such as (imputed) genotypes and (exome) sequencing data. The imputed genotype data provides two quality metrics, which are INFO score and imputed alleles probability. The quality metrics can be used for filtering imputation results per individual variant. We determined the optimal imputation quality metrics in order to extract only the most genuine LoF variants.  We used whole exome sequencing (WES) data from UK biobank (UKBB) as a gold standard for evaluating the optimal quality metrics to obtain the most genuine LoF variants from imputed genotypes data. We extracted WES and genotypes data of 4,476 randomly selected UKBB participants. Genotypes were phased and imputed using SHAPEIT2 and IMPUTE2, respectively (Howie et al. 2009; Howie et al. 2012; Delaneau et al. 2011). Phasing and imputation were performed using a combined reference panel from the 1000 Genome project phase 3 (1000 Genomes Project Consortium et al. 2015) and Genome of the Netherlands (GoNL) study (Boomsma et al. 2014). The imputed genotypes were submitted to LoFTK using a range of info scores and imputed allele probabilities in order to obtain various numbers of LoF variants for each subset. On the other hand, the UKBB provided unphased WES data, thus we phased the data using SHAPEIT2 and then converted to VCF. We retrieved overlapped variants between phased exomes and imputed genotypes data (Supplementary Figure 1).

In the imputed genotype data, we analysed LoF variants for three different datasets, where the first has INFO > 0.3, the second has INFO > 0.6 and the third has INFO > 0.9. We compared the existence of each LoF variant between each imputed genotypes dataset and WES with considering the imputed allele probabilities (0.01 – 0.1) for that variant (Supplementary Figure 1). For each individual, each LoF variant from the three imputed datasets was matched to WES, in order to count the false negative (average of LoF variants found in WES data but not in imputed data) and false positive (average of LoF variants found in imputed data but not in WES data) (Supplementary Table 1). The imputed dataset with INFO > 0.9 shows an optimal prediction of true LoF variants, because it has less false positive 2-copy LoF variants (~ three variants) compared to the others (0.1 and 0.3). However, selecting an optimal imputed allele probability was difficult due to the lack of apparent variations.

### Compound heterozygous LoF variants

Compound heterozygous (CH) variants occur when both parents donate a single LoF allele to proband at distinct loci within the same gene (Kamphans et al. 2013). LoFTK can annotate CH LoF variants, which introduce 2 inactive copies of genes. We used trio-families data from the Genome of the Netherlands (GoNL) study (Illumina Immunochip microarray SNP data) (Boomsma et al. 2014) to validate the transmitting of real CH LoF variants from parents to probands. We included 760 samples from the GoNL cohort, which they are family-based trios. We excluded variants with a call rate ≤ 0.99, Hardy-Weinburg equilibrium (HWE) ≤ 0.001 and monomorphic variants. After data filtrations, we used TOPMed imputation server to impute missing genotypes (Taliun et al. 2021). As a post-imputation quality control step, we excluded variants with imputation quality score < 0.3 (INFO < 0.3) and monomorphic variants.

We utilized LoFTK to predict LoF variants and genes in the imputed genotypes of 250 families. We discovered 250 CH LoF variants generating 2-copy LoF genes in 164 probands. There were 240 (96%) genuine transmissions of CH LoF in parent-offspring with 10 false transmissions.

### Supplementary table

| Imputed allele probabilities | info score | | | | | |
|---|---|---|---|---|---|---|
| | 0.3 | | 0.6 | | **0.9** | |
| | False negative | False positive | False negative | False positive | False negative | False positive |
| 0.01 | 21.69 | 19.59 | 22.18 | 19.24 | 23.63 | 16.42 |
| 0.02 | 21.40 | 19.78 | 21.91 | 19.38 | 23.42 | 16.47 |
| 0.03 | 21.23 | 19.90 | 21.75 | 19.48 | 23.30 | 16.49 |
| 0.04 | 21.10 | 19.99 | 21.64 | 19.54 | 23.22 | 16.51 |
| 0.05 | 21.01 | 20.06 | 21.55 | 19.59 | 23.16 | 16.51 |
| 0.06 | 20.90 | 20.14 | 21.46 | 19.64 | 23.09 | 16.52 |
| 0.07 | 20.81 | 20.21 | 21.38 | 19.69 | 23.04 | 16.54 |
| 0.08 | 20.74 | 20.27 | 21.33 | 19.74 | 23.00 | 16.55 |
| 0.09 | 20.68 | 20.33 | 21.27 | 19.79 | 22.96 | 16.56 |
| 0.1 | 20.61 | 20.40 | 21.22 | 19.84 | **22.92** | **16.58** |

Supplementary Table 1: False positive and false negative values from matching the LoF variants between exome and subgrouped imputed genotypes data for each individual.

### Supplementary Figure

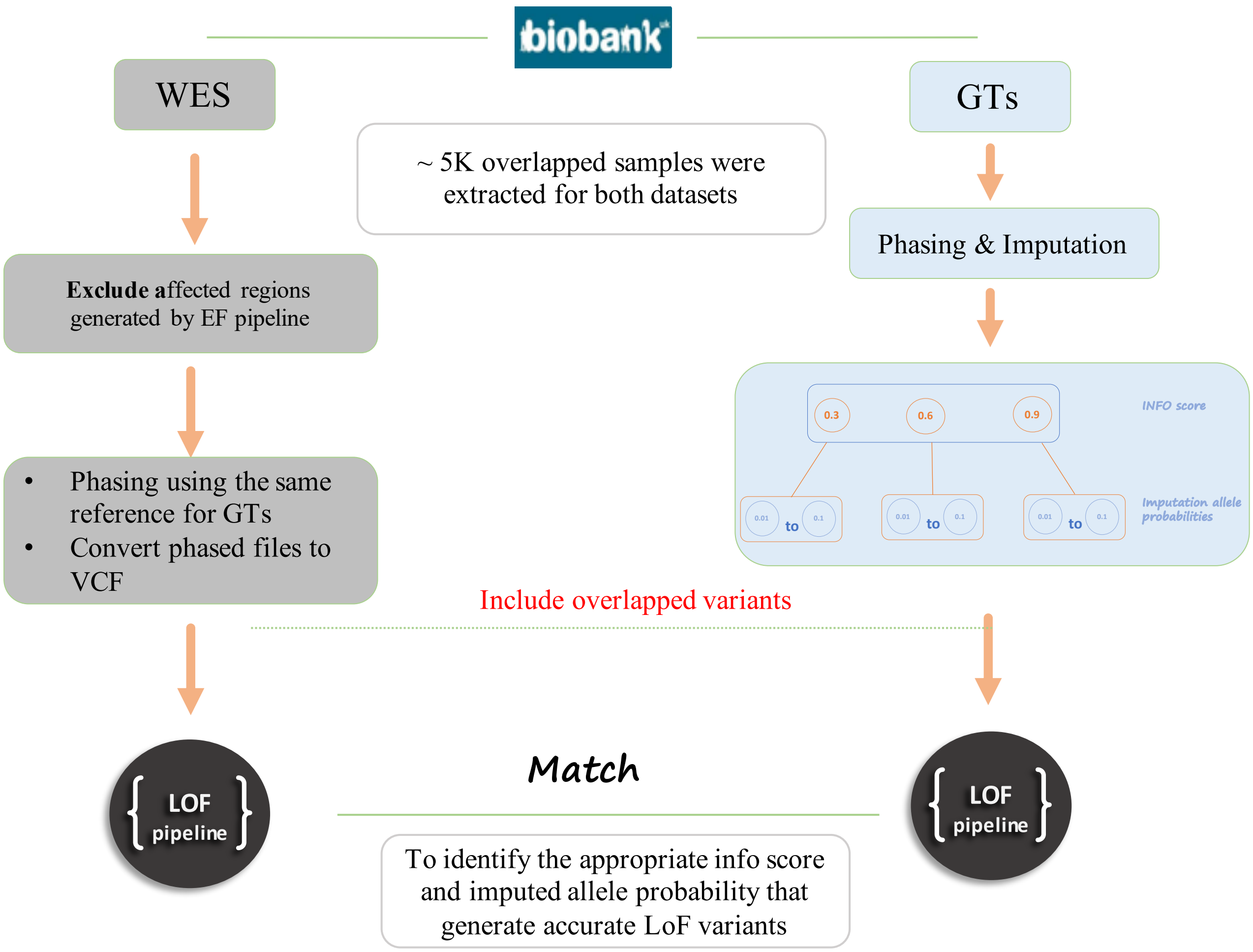


Supplementary Figure 1: The workflow for achieving the optimal imputation quality measure

### References


1000 Genomes Project Consortium, Adam Auton, Lisa D. Brooks, Richard M. Durbin, Erik P. Garrison, Hyun Min Kang, Jan O. Korbel, et al. 2015. "A Global Reference for Human Genetic Variation." *Nature* 526 (7571): 68–74. https://doi.org/10.1038/nature15393.

Boomsma, Dorret I., Cisca Wijmenga, Eline P. Slagboom, Morris A. Swertz, Lennart C. Karssen, Abdel Abdellaoui, Kai Ye, et al. 2014. "The Genome of the Netherlands: Design, and Project Goals." *European Journal of Human Genetics: EJHG* 22 (2): 221–27. https://doi.org/10.1038/ejhg.2013.118.

Delaneau, Olivier, Jonathan Marchini, and Jean-François Zagury. 2011. "A Linear Complexity Phasing Method for Thousands of Genomes." *Nature Methods* 9 (2): 179–81. https://doi.org/10.1038/nmeth.1785.

Howie, Bryan, Christian Fuchsberger, Matthew Stephens, Jonathan Marchini, and Gonçalo R. Abecasis. 2012. "Fast and Accurate Genotype Imputation in Genome-Wide Association Studies through Pre-Phasing." *Nature Genetics* 44 (8): 955–59. https://doi.org/10.1038/ng.2354.

Howie, Bryan N., Peter Donnelly, and Jonathan Marchini. 2009. "A Flexible and Accurate Genotype Imputation Method for the next Generation of Genome-Wide Association Studies." *PLoS Genetics* 5 (6): e1000529. https://doi.org/10.1371/journal.pgen.1000529.

Kamphans, Tom, Peggy Sabri, Na Zhu, Verena Heinrich, Stefan Mundlos, Peter N. Robinson, Dmitri Parkhomchuk, and Peter M. Krawitz. 2013. "Filtering for Compound Heterozygous Sequence Variants in Non-Consanguineous Pedigrees." PloS One 8 (8): e70151.

Taliun, Daniel, Daniel N. Harris, Michael D. Kessler, Jedidiah Carlson, Zachary A. Szpiech, Raul Torres, Sarah A. Gagliano Taliun, et al. 2021. "Sequencing of 53,831 Diverse Genomes from the NHLBI TOPMed Program." Nature 590 (7845): 290–99.
